# Human Cytomegalovirus UL4 is Required for Viral Reactivation via Cellular Reprogramming in Latently Infected CD34+ Progenitor Cells and Humanized Mice

**DOI:** 10.64898/2026.06.28.735103

**Authors:** Chris Held, Kamryn Pfenning, Lindsey B. Crawford

## Abstract

Human cytomegalovirus (HCMV) remains a significant cause of morbidity and mortality after both solid organ and hematopoietic stem cell transplant, due to bone marrow stem cell engraftment failure, myelosuppression, and immunosuppression. While direct infection and viral replication lead to disease, latent infection in the CD34+ hematopoietic progenitor cell (HPC) pool has direct and indirect effects on hematopoiesis. Studies from several laboratories support the hypothesis that HCMV latency and reactivation are intrinsically linked with the state of the cell. We, and others, have previously demonstrated roles for different viral gene products in regulating both cellular differentiation and the balance between viral latency and reactivation, including the viral RL11 proteins UL7 and UL8. In this study, we show that UL4 is expressed during latency and required for reactivation in HPCs, THP-1 monocytes, and humanized mice. describe both a cell-specific and viral lifecycle-specific role for the RL11 gene UL4 during HCMV infection. Additionally, we demonstrate that UL4 plays specific roles in controlling essential cellular functions including aspects of proliferation and differentiation and immune signaling to assist in establishing a virus-favorable environment in progenitor cells. This study identifies a novel viral reactivation factor that has implications for both viral control and alteration of hematopoiesis in transplant patients.

## Introduction

Pathogenesis following human cytomegalovirus (HCMV) infection *in vivo* results from (1) viral latency-induced myelosuppression and/or (2) acute CMV disease following reactivation which lead to severe disease in hematopoietic and solid organ transplant patients(1–6). While previous work, both ours and others, have defined the roles of some viral and cellular genes (as we recently reviewed(7)) in controlling latency and reactivation, a full mechanistic understanding of HCMV latency and reactivation is still poorly defined.

Herpesvirus latency in general, and HCMV latency specifically, is defined as the ability of the virus to enter a cell and maintain the viral genome, without producing infectious virus. While infectious virus can be recovered from primary peripheral blood monocytes from seropositive patients through allogeneic stimulation *ex vivo*, infection of monocytes *in vitro* results in limited expression of immediate early (IE) gene products and no infectious virus production(8–11). CD34^+^ hematopoietic progenitor cells (HPCs), provide a critical reservoir of latent HCMV(12, 13) and infection of HPCs contributes to the hematopoietic abnormalities observed in transplant patients(14–17). *In vivo*, latently infected HPCs exit the bone marrow in response to cytokine/growth factor signaling, traffic to the periphery and differentiate into monocytes and tissue macrophages(14, 18, 19). These differentiation processes provide a cellular environment appropriate for viral replication and reactivation(18, 20, 21). In parallel, direct infection of monocytes promotes differentiation to macrophages(18, 21–23) and infection of either HPCs or monocytes specifically alters their differentiation(24–28). The current working hypothesis is that HPCs provide the latent reservoir, monocytes disseminate the virus, and macrophages produce infectious virus for spread throughout the host.

Previous work demonstrated that two genes in the HCMV RL11 region are key to latency establishment, maintenance, and reactivation. The HCMV RL11 region is a multigene family composed of 12 open reading frames. Eleven are highly conserved between different CMV species and most are predicted to encode glycoproteins with some homology to Adenovirus E3 proteins(29). None of the RL11 proteins are required for lytic replication in fibroblasts(30–32). We previously described a role for UL7, a secreted RL11 glycoprotein, both in differentiation of CD34^+^ HPCs and monocytes as well as for efficient reactivativation both *in vitro* and *in vivo* in humanized mice(25). Later collaborative work identified UL7 as a co-regulator of cellular apopotsis(33) and the co-expressed RL11 protein, UL8, as an additional regulator of viral reactivation(27).

UL4 is a member of the RL11 gene family that encodes a 150 amino acid protein. Previous studies first identified UL4 as an early glycoprotein in 1989(34) and while it is highly homologous between HCMV strains and conserved between primate species suggesting a critical function, little more has been studied. During lytic replication, multiple UL4 transcripts have been identified, three with immediate-early and one with late kinetics(35). Studies using large deletions of the RL11 region also demonstrate that the region including UL4 is not essential for viral replication in fibroblasts(36, 37). However, in progenitors and myeloid lineage cells, the expression kinetics and role of UL4 is less clear. In this study, we examined the functional role of UL4 during HCMV latency and reactivation in multiple models including CD34+ HPCs, THP-1 monocytes, and in vivo in humanized mice. We observed that virus lacking UL4 protein is unable to reactivate both in vitro and in vivo. Prior work from the Stanton lab demonstrated that UL4 is a secreted protein that inhibits TRAIL-mediated apoptosis in NK cells(38). We also found that UL4 expression in hematopoietic cells has specific effects on hematopoiesis, including some regulation of immune signaling, indicating that UL4 is an important factor for HCMV reactivation through regulation of cellular functions including differentiation.

## Results

### UL4 is not required for viral replication in fibroblasts

Since prior studies demonstrating that the RL11 region is not required for viral replication in fibroblasts used large deletions of multiple genes(36, 37), but several viral gene regions with significance for latency and reactivation have been shown to have co-regulation of functional outcomes and/or specific co-expression (e.g., multiple isoforms of UL136(39, 40) or UL7/UL8(27)), we first generated a stop mutant of UL4 in the TB40/E-GFP BAC lacking expression of UL4 protein following viral reconstitution. HCMV-UL4stop shows no defect in viral reconstitution (data not shown) and no change in viral replication in fibroblasts (**Figure 1**) either in cell-associated (**Figure 1A**) or secreted virus (**Figure 1B**) as measured by infectious titer. The UL4 mutation is maintained with at minimum 2 out of 3 stop mutations through at least passage 7 (data not shown).

**Figure 1.**
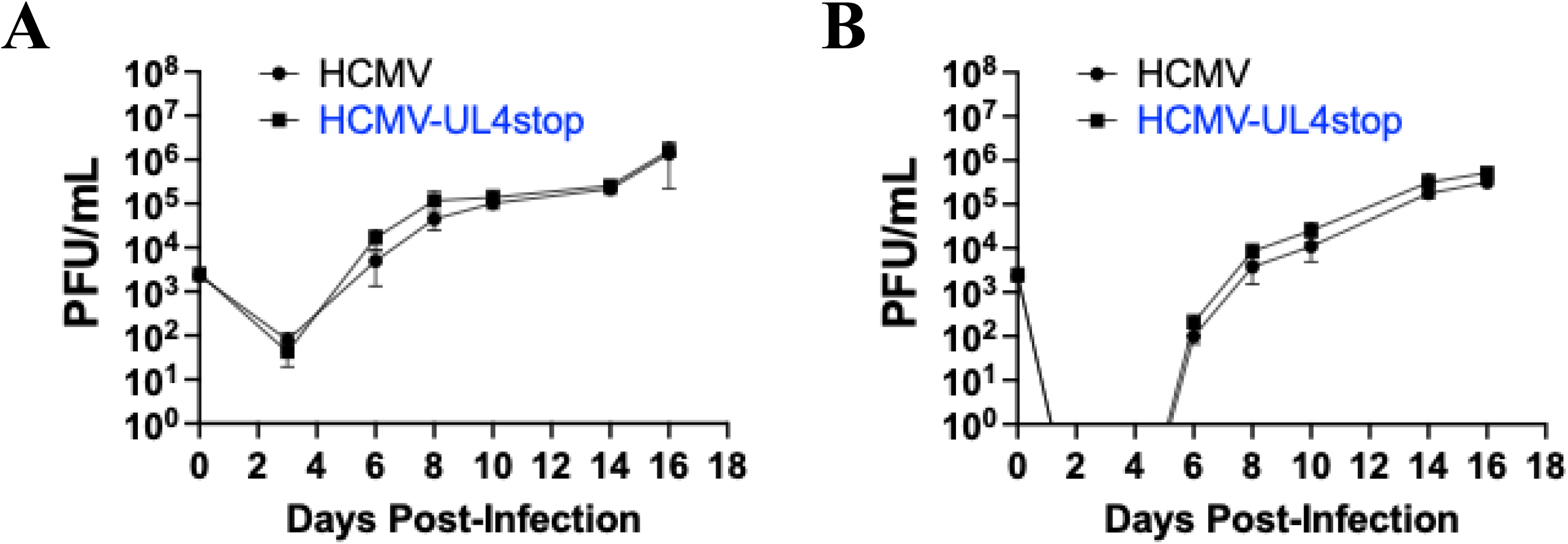
UL4 is not required for replication during lytic infection in fibroblasts. NHDFs were infected with HCMV or HCMV-UL4stop at an MOI of 0.02 for a multi-step growth curve. Cells (A) and supernatant (B) were collected at the indicated times and stored at −80. Viral titer was determined by plaque assay on NHDFs following a 2 week incubation. Data shown is average of two replicate titers per experimental as average and standard error of the mean for three independent experiments.

### UL4 is expressed as a glycosylated protein and during latency in hematopoietic cells

During lytic replication, multiple UL4 transcripts have been identified, three with immediate-early and one with late kinetics(35). In primary hematopoietic cells, transcriptional analysis detects UL4 at 6dpi and in clinical samples(41). Other analysis using single cell sequencing assessing hematopoietic lineage cellular subpopulations(42), however did not separate UL4 from UL5, leaving the expression pattern, and therefore insights into the role of UL4 undefined. Reanalysis of published data from the Goodrum lab using THP-1 monocytes in an established latency model indicates that UL4 is expressed during THP-1-latency (**Supplemental Figure 1**). We also confirmed that UL4 can be expressed using common plasmid-based expression systems (Figure 2A) for stable protein production and characterized UL4 expression during latency and reactivation in CD34+ HPCs. Briefly, CD34^+^ HPC were infected with a HCMV that expresses GFP (strain TB40/E) for 2d, purified by fluorescence-activated cell sorting (FACS), and seeded into long-term bone marrow culture (LTBMC) over stromal cell support as previously described(43). Cells were harvested during latency establishment and maintenance at the indicated times, RNA extracted, and cDNA generated for qRT-PCR analysis of UL4 at initial plating (2dpi), latency establishment (6 and 8dpi), latency maintenance (12dpi), and immediately before reactivation (14dpi) (**Figure 2B**). These data indicate that UL4 is one of a limited number of viral gene products expressed during latency.

**Figure 2.**
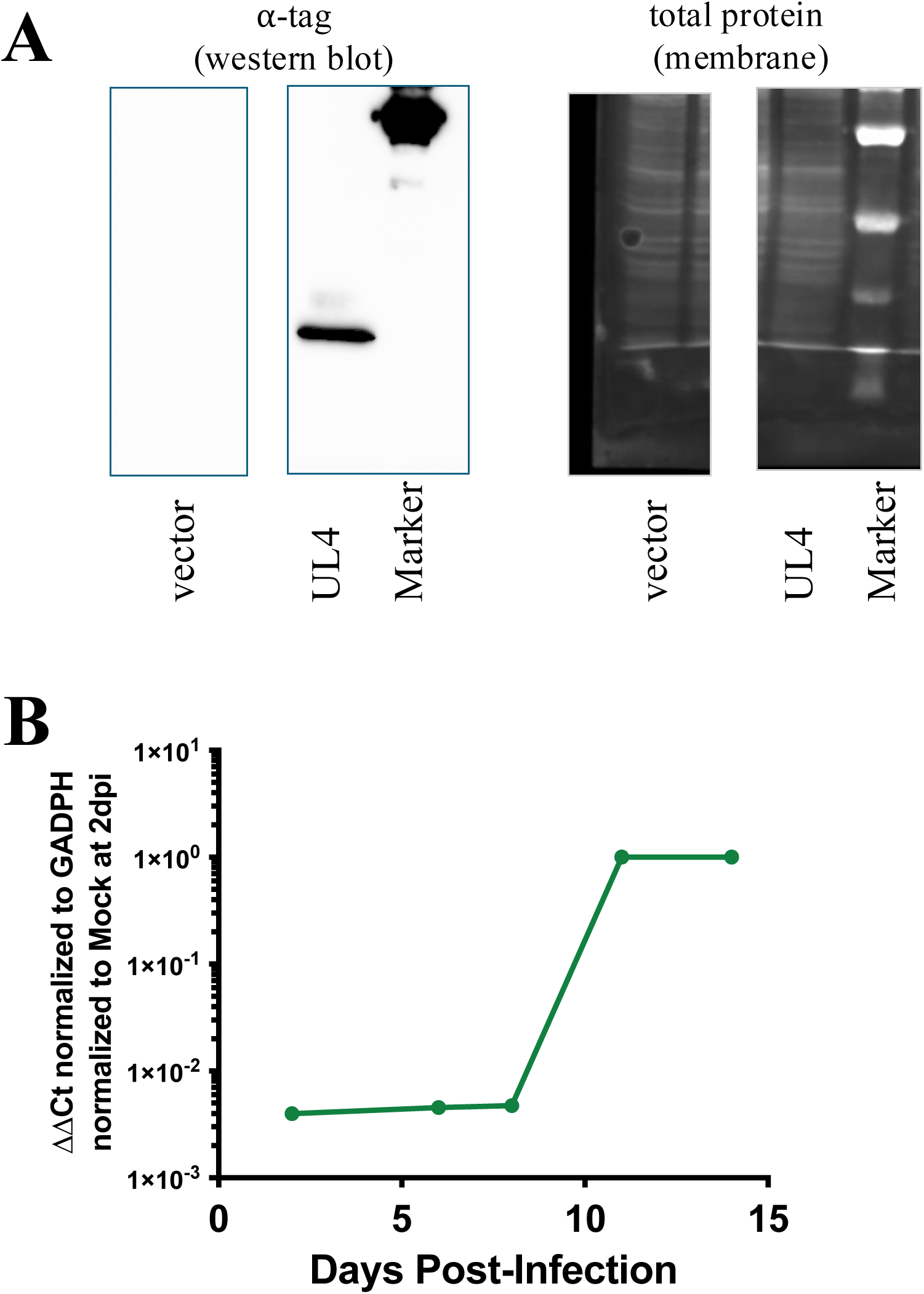
UL4 is expressed late during HPC infection. A) 293T cells were transfected with a plasmid vector expressing UL4 or empty parent plasmid using Lipofectamine 2000. Cells were harvested in cell lysis buffer (Cell Signaling Technologies) at 48h. Total protein was normalized by Bradford Assay and run on an 8% SDS-PAGE gel with sample buffer. Total protein was stained in the transferred membrane with the Licor Total Protein Stain prior to antibody detection of the protein tag. **B)** CD34^+^ HPCs were infected with HCMV for 2d, FACS isolated for a pure population of viable CD34^+^GFP^+^ HPCs, then cultured with stromal cell support for an additional 12d to establish latency as previously described. RNA was extracted at the indicted times post-infection (2, 6, 8, 10, 14dpi) and copies of HCMV UL4 and human GADPH determined by qRT-PCR.

### UL4 is required for viral reactivation

As limited viral gene products are expressed during latency and prior data indicates of those currently studied, that viral genes expressed during latency play a role in latency and/or reactivation, we analyzed whether UL4 is required for reactivation. Briefly, as previously described(43), CD34+ HPCs were infected and cultured as above with either wild-type HCMV or HCMV-UL4stop. After 12 days of culture in LTBMC, latently infected HPCs were seeded onto monolayers of permissive fibroblasts in cytokine-rich media to promote myeloid differentiation. The HPC/fibroblast co-cultures were analyzed weekly by limiting dilution assay for the fraction of GFP+ wells for up to four weeks to determine the frequency of infectious centers. As shown in **Figure 3A**, loss of UL4 results in an inability of the virus to reactivate. Analysis of viral DNA in both wild-type HCMV and HCMV-UL4stop infected CD34+ HPCs at 14dpi (12days latent) indicated no significant change in HCMV genomes between viruses across multiple HPC donors (**Figure 3B**). The requirement of UL4 for viral reactivation is also maintained under cellular selection pressure, including in primary donors that do not support latency establishment (**Supplemental Figure 3A** and **B**) or following expansion of HPCs prior to infection (**Supplemental Figure 3C**). These results indicate that the inability of the UL4stop mutant to reactivate is not due to the loss of viral genomes during latency.

**Figure 3.**
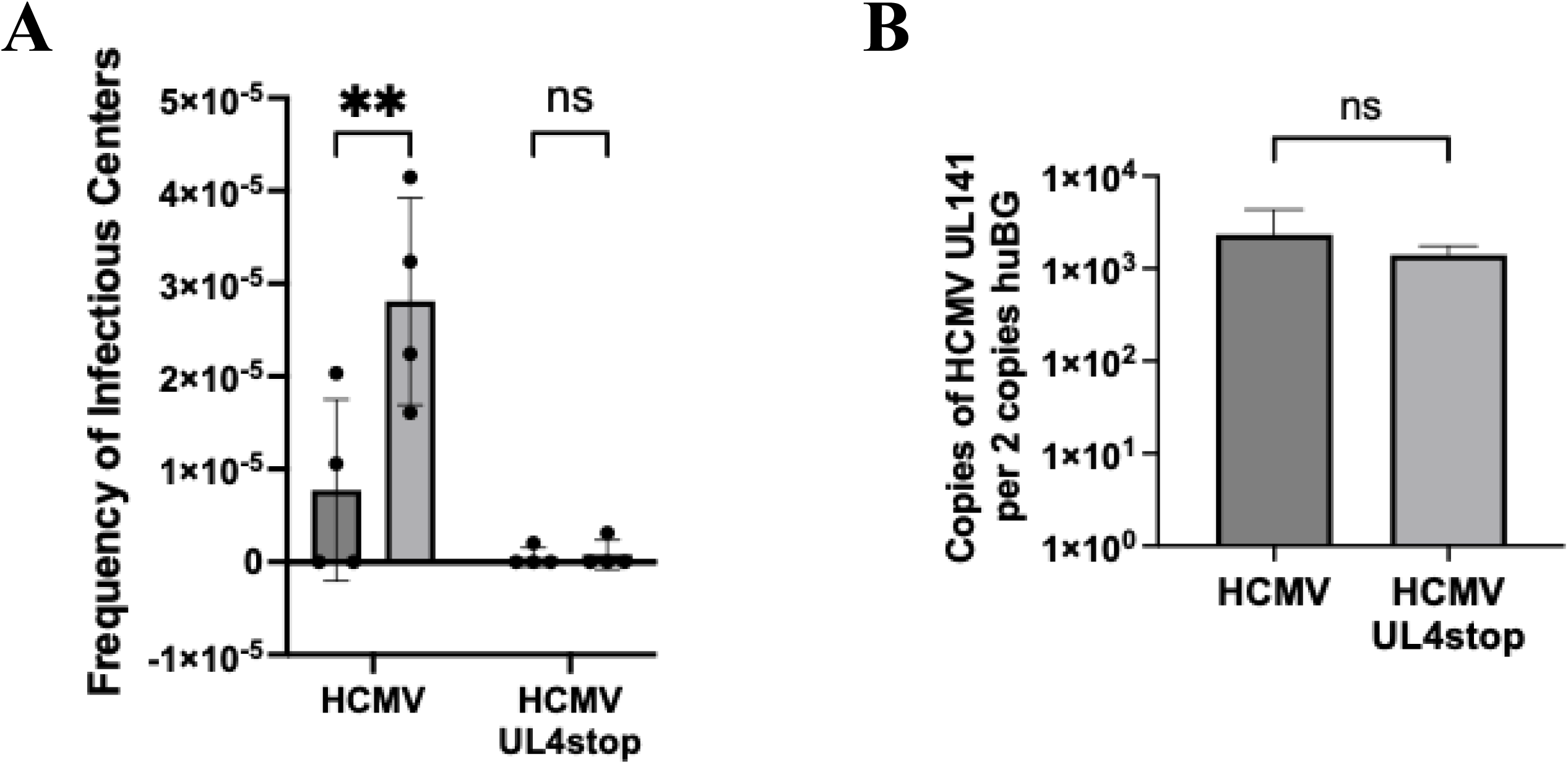
UL4 is required for latency and reactivation in CD34+ HPCs. **A)** CD34+ HPCs infected with HCMV or HCMV lacking UL4 (HCMV-UL4stop), were sorted, and cultured for latency as in Figure 2. At 14dpi (12d latent), equivalent populations of cells were either directly plated on fibroblasts in the presence of cytokine support (reactivation) or lysed and plated on fibroblasts to measure the amount of virus present during latency (pre-reactivation), and the frequency of infectious centers calculated by ELDA. Data shown is the absolute frequency of infectious center formation for 4 independent primary HPC donors. Individual donors are shown separately in Supplemental Figure 3. **B)** Total DNA was extracted from a fraction of the cells from donors 3 (Supplemental Figure 3C) and 4 (Supplemental Figure 3D) at 14dpi (12d) latent. Viral genomes were determined on a per cell basis using primers and probes for HCMV UL141 and human beta-globin as previously described. Data shown is the mean +/- standard error of the mean for these donors.

As HCMV has previously been shown to have cell-specific effects, especially in latency and/or persistence establishment in cells of different lineages and/or differentiation stages (7), we also examined the ability of HCMV-UL4stop to reactive in an established model of monocyte latency using the THP-1 cell line. Here, cells are infected with either wild-type or HCMV-UL4stop for 24, washed with trypsin and PBS, then plated in fresh media. To assess the role of UL4 during infection in THP-1 cells, we then collected cell fractions and cell-free supernatant at the indicated times in parallel with cells for RNA extraction. THP-1 have previously been shown to establish a monocyte-like latency phenotype in 4-5dpi and viral reactivation or recovery can be induced by plating cells treatment with the phorbol ester, TPA(44, 45). As shown in Figure 4 UL4 is required for efficient replication of HCMV in THP-1 cells following TPA treatment at 5dpi (**Figure 4A**) and for release of infectious virus into the supernatant (**Figure 4B**). As latency and reactivation in THP-1 cells are often described via regulation of viral gene classes (e.g., IE1 and UL138)(44–51) and to assess the role of UL4 on early viral gene expression, we also assessed whether the loss of UL4 alters immediate early (as measured by IE1 expression, **Figure 4C**) or latency (measured by UL138, **Figure 4D**) gene expression. Interestingly we found that both IE1 and UL138 expression is decreased during HCMV-UL4stop infection.

**Figure 4.**
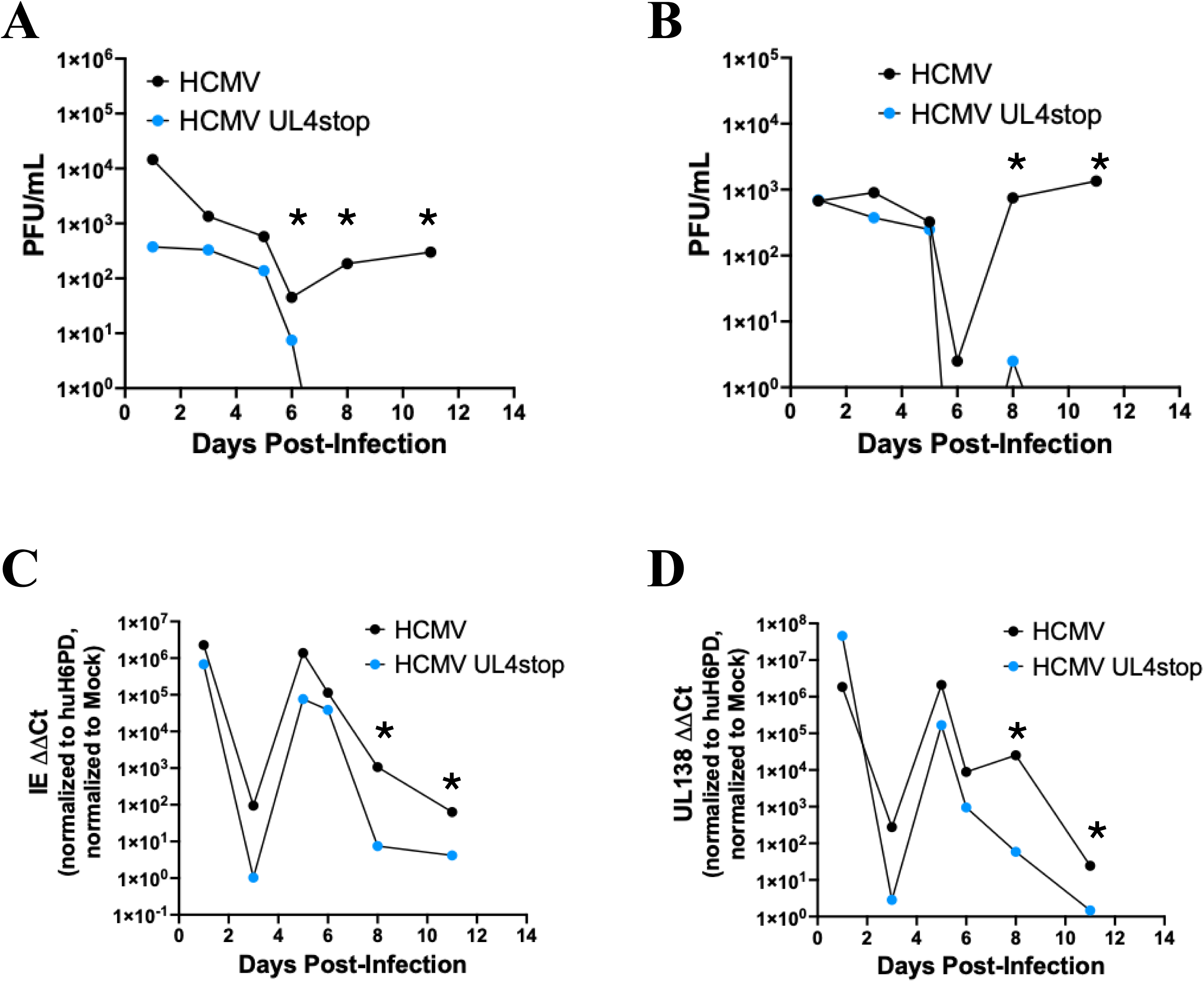
UL4 is required for latency and reactivation in THP-1 cells. **A)** THP-1 monocytes were infected as previously described and cells and supernatants collected daily. Viral reactivation was induced by TPA treatment at 5dpi with cells and supernatants also collected at 6, 8, 11dpi (=1, 3, 5d post-reactivation). Data shown is representative of 3 independent experiments. **B and C)** In parallel, RNA was extracted from cells using TRIZOL, cDNA synthesized and qRT-PCR performed for HCMV IE1 (B) and HCMV UL138 (C) as normalized to human H6PD.

To further extend the results of Figures 3 and 4, we also examined the ability of HCMV-UL4stop to reactivate in a humanized mouse model of viral latency and reactivation. In this model NODscid-IL2Rγc null mice (NSG) are engrafted with human CD34+ HPCs (huNSG), a commonly used platform to study human-tropic viruses. As previously described, huNSG mice infected with HCMV establish latency and permit viral reactivation following treatment with G-CSF(52). HCMV-infected huNSG mice have previously been valuable in demonstrating unique roles of viral latency and reactivation across viral strains(53) and for specific gene products including UL7(25), UL8(27), US28(24), and the UL135/UL138 locus(40, 54), and more expanded huNSG models for specific viral immune responses(55). As shown in **Figure 5**, both wild-type HCMV and HCMV-UL4stop establish infection in hematopoietic organs [spleen (**Figure 5A**) and liver (**Figure 5B**)], however only wild-type HCMV and not HCMV-UL4stop reactivated in these tissues. Similar to in vitro analysis in CD34+ HPCs (**Figure 3B**), HCMV-UL4stop does not show a loss of viral genomes in vivo (Figure 5A and **B**) indicating that the viral genome is maintained but there is a defect in reactivation.

**Figure 5.**
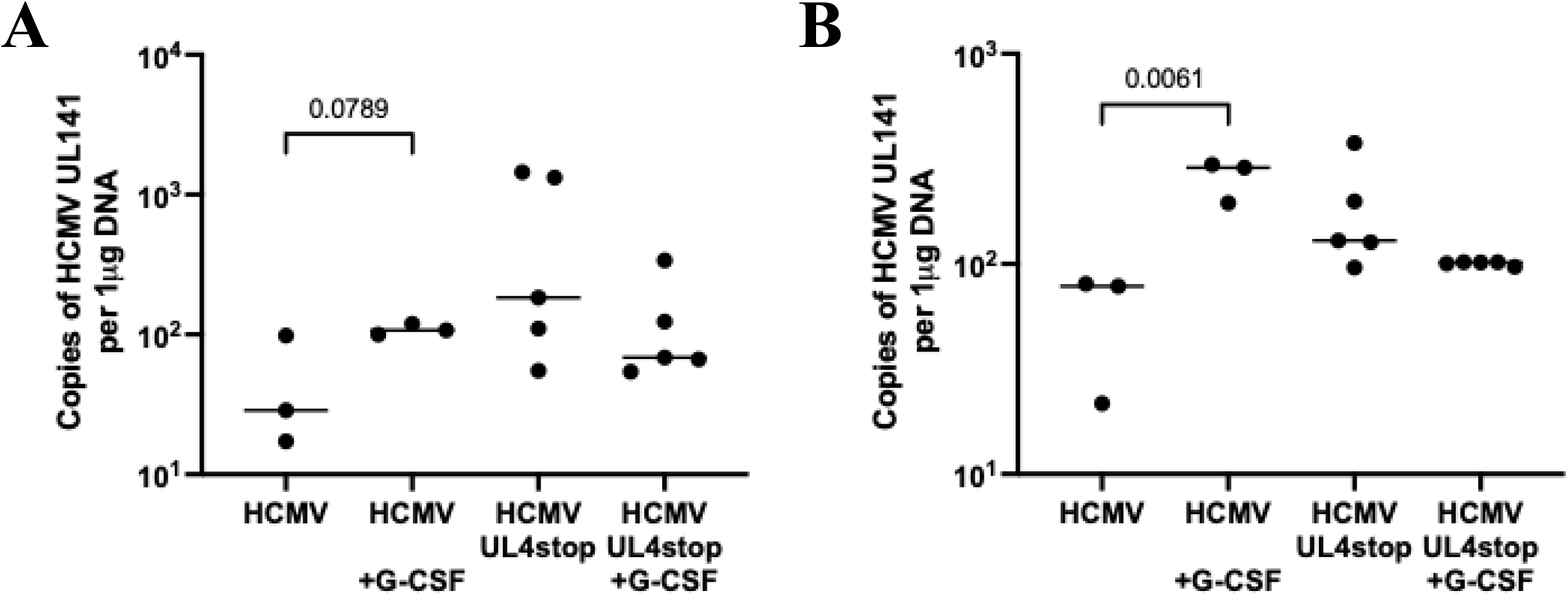
UL4 is required for reactivation in vivo in humanized mice. Humanized mice (N=3-5/group) were generated and infected with HCMV as previously described. After 8 weeks, reactivation was induced by G-CSF treatment for 7 days and viral genomes measured in the spleen (**A**) and liver (**B**) by qPCR. Data shown is average and standard error of the mean for two slices of spleen and four slices of liver per humanized mouse, each mouse represented by one data point. Significance determined by comparison of latent vs reactivated for each virus and reactivated wild-type vs UL4stop. Significant groups are listed by p values in the figure, all other groups are not significantly different.

### UL4 directly induces myelopoiesis when expressed endogenously

Since HCMV reactivation has previously been linked to cellular differentiation including globally in transplant donors(56), humanized mice(52, 57), and as we have previously shown for another RL11 gene, UL7(25). To determine if UL4 alone is sufficient to induce myeloid differentiation, we overexpressed UL4 in CD34+ HPCs and assessed myeloid colony formation as previously described(24, 25, 58). We compared cells transduced with an adenovirus without insert (Ad-empty) or GFP alone (Ad-GFP) with Ad-UL4. As shown in **Figure 6A**, expression of UL4 substantially increased the ability CD34+ HPCs to form myeloid colonies as assessed by total colony formation. To determine if this effect is due to specific changes in lineage differentiation or a more global proliferative effect during the differentiation assay, we assessed both lineage differences (**Figure 6B**) and HPC proliferation (**Figure 6C**). Both myeloid (CFU-GM) and erythroid (BFU-E) colonies are increased in the presence of UL4 (**Figure 6B**) and a modest increase in total HPC proliferation is observed in the absence of differentiation stimulus (**Figure 6C**).

**Figure 6.**
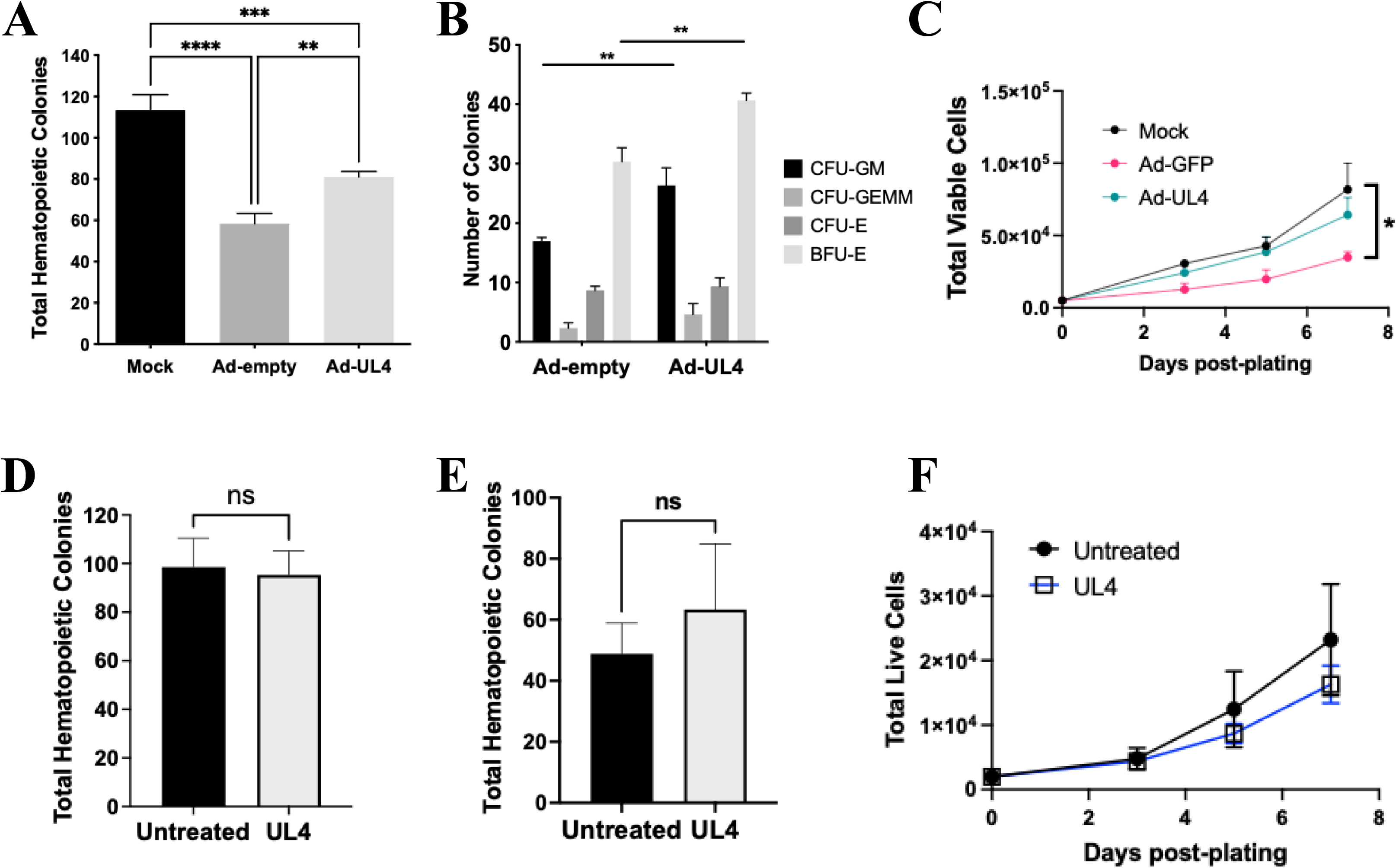
Endogenous but not exogenous UL4 controls HPC functional outcomes. HPCs infected with adenovirus expressing UL4 or control and plated in semi-solid media enriched with cytokines (Methocult) for differentiation of single HPCs to myeloid or erythroid lineages (**A, B**) or in cytokine enhanced liquid media for proliferation **(C)**. Or HPCs were treated with exogenous UL4 protein and plated in semi-solid media enriched with cytokines (Methocult) for differentiation of single HPCs to myeloid or erythroid lineages (**D, E**) or in cytokine enhanced liquid media for proliferation **(F)**. Data shown in A, B, C, and D are representative of 3 independent experiments. Additional data is shown in Supplemental Figure 4. Data shown in D and E is average and standard error of the mean for 3 independent experiments shown combined. Myeloid colonies **(A, B, D, E)** were enumerated at 14d post-plating using standard morphological characteristics. Proliferation (C, F)) was determined by total viable cell counts using Trypan Blue at 3, 5, and 7 days post-plating. *p<0.05, NS = not significant.

As there is a previously demonstrated key role for ligand signaling in the connection between viral reactivation and cellular differentiation both through viral receptors (e.g. US28(24)) or through viral-encoded ligands signaling through cellular receptors (HCMV UL7 through cellular FLT3(25)) and prior work from the Stanton lab demonstrated that UL4 is a secreted factor that may act as a signaling ligand in NK cells (38), we asked whether UL4 signals as a ligand in HPCs. To determine if exogenously expressed UL4 induces HPC differentiation, we used purified pUL4 added to HPC cultures during myeloid colony formation. Interestingly, we found that exogenous pUL4 has no effect on myeloid colony formation (**Figure 6D**) from either of two independently, commercially generated pUL4 batches (Figure 6D and **E**). We also found that pUL4 has no significant effect on HPC proliferation.

We therefore hypothesized that endogenously expressed UL4 may play a role in hematopoietic cell signaling either directly or induction of secreted factors to regulate the hematopoietic environment. Preliminary analysis demonstrates that while loss of UL4 does not significantly alter the transcriptome early in lytic infection in fibroblasts (data not shown), overexpression of UL4 alone in HPCs alters a subset of select hematopoietic-related genes similarly to HCMV infection (data not shown). As several of these factors converge on the a nuclear-related signaling network related to immune signaling, and we have previously shown a key role of the master cellular cytokine TGFβ in regulating myelopoiesis linked to viral latency(58) and secreted UL4 has been shown to inhibit NK cell degranulation by binding to TRAIL(38), we analyzed whether UL4 alone induces immune signaling in the presence of viral stimulation. Using the adenovirus expression system described above, HPCs were infected with Ad-GFP (control) or Ad-UL4 for 2 days, washed, and replated in fresh cytokine-enriched media for 24h and supernatants collected. To assess for global effects, viral transduction was controlled, but bulk populations of transduced/expressing and bystander HPCs were analyzed using multi-factor cytokine analysis was performed using LEGENDplex assays. As shown in **Figure 7**, both MCP-1 (**Figure 7A**) and IL-8 (**Figure 7B**) are preferentially upregulated in the secreted environment of UL4-expressing HPCs. However, these changes are insufficient to result in myeloid differentiation, as bulk populations of the same HPCs show no significant change in myeloid colony formation (**Figure 7C**). Combined, this suggests that UL4 has specific effects in HPCs directly expressing endogenous UL4, but that neither exogenous pUL4 nor UL4-regulated cytokine changes are sufficient to cause global differentiation changes.

**Figure 7.**
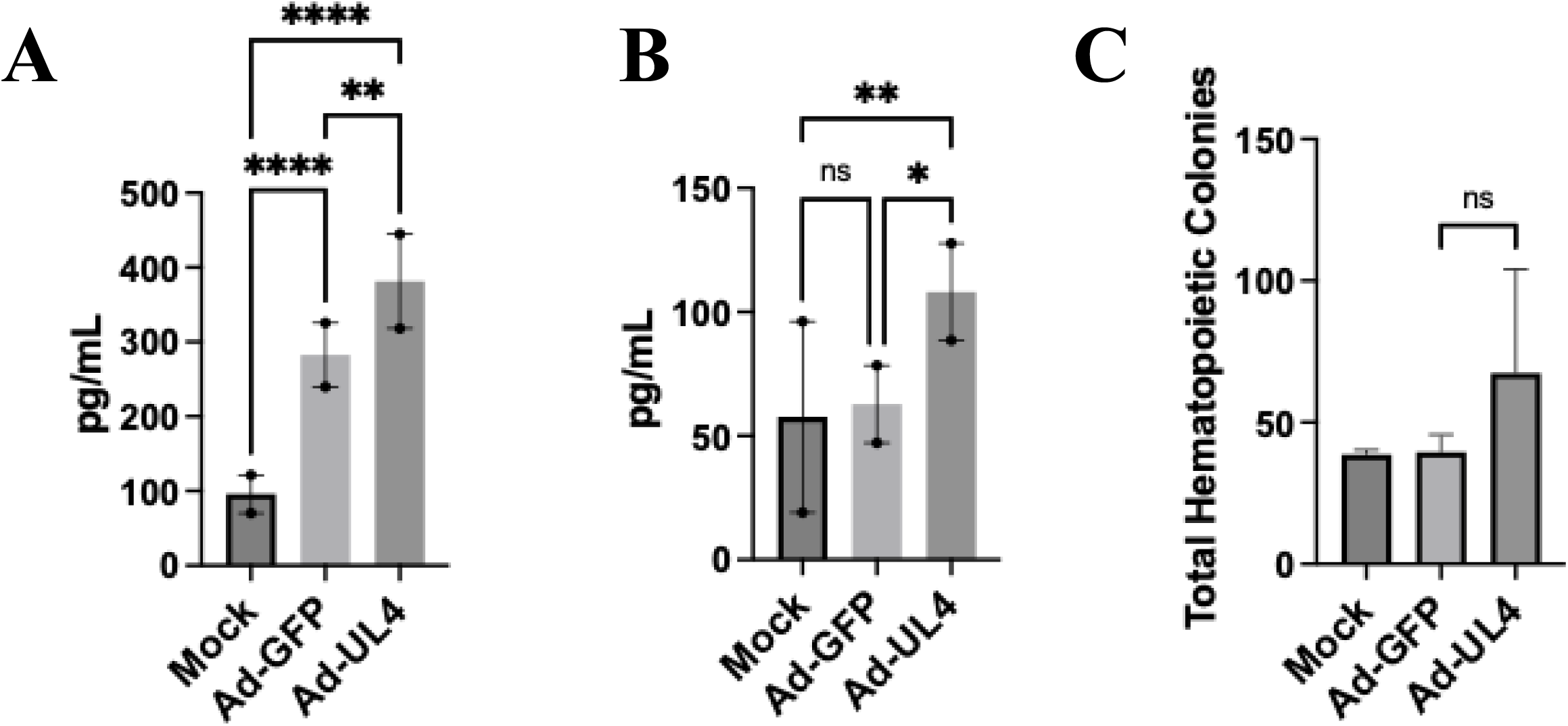
Endogenous UL4 upregulates environmental cytokines without a global effect on myeloid differentiation. HPCs were infected with adenovirus expressing UL4 or control for 2 days, washed, and plated in fresh media for 24h. Cell-free supernatant was harvested and stored frozen prior to analysis with LegendPlex multi-plex cytokine assays (Biolegend). Data shown is the average of technical duplicates for two independent experiments using different donors. Data shown is for cytokines above the limit of detection of the assay for Mock-infected HPCs and that demonstrate a change following viral transduction for MCP-1 (**A**) and IL-8 (**B**). **C)** In parallel bulk HPCs were plated at 2d in semi-solid media enriched with cytokines (Methocult) for differentiation of single HPCs to myeloid or erythroid lineages. Total colonies were enumerated at 14d post-plating using standard morphological characteristics. *p<0.05, NS = not significant.

## Discussion

In summary, we identify UL4 as a previously unrecognized regulator of HCMV reactivation. We demonstrate that UL4 is dispensable for productive lytic replication in fibroblasts (**Figure 1**) but is both expressed during latency in HPCs (**Figure 2B**) and required for reactivation in HPCs (**Figure 3A**), THP-1 cells (Figure 4A and **B**), and in vivo in humanized mice (Figure 5A and **B**). Importantly, the reactivation defect occurs despite viral genome maintenance (**Figure 3B** and Figure 5A and **B**), indicating UL4 is required for reactivation or exit from latency but not latency establishment or maintenance. These findings place UL4 in the small subset of viral latency and reactivation regulators including US28, UL135/UL138 locus, UL7 and UL8. The overlap between UL4’s requirement in the latent reservoir of HPCs, the persistence reservoir of THP-1s and the spectrum of HCMV biology in vivo suggests that UL4 contributes to a conserved mechanism regulating reactivation across cell and model system differences.

The reduced expression of both IE1 and UL138 during infection with the UL4stop virus (Figure 4C and **D**) suggests that UL4 may function as a regulator of viral transcription either directly or indirectly. It is well established that IE1 gene expression is differentially regulated during HCMV infection, including downregulation during latency in comparison to “classic latency factors” such as US28 and UL138. However, it is also know that these effects are cell-type specific (48, 59, 60) and, outside of US28 (61) and UL138 (62), no other viral factors have been identified as central to this phenotype. Additionally, suppression of IE expression can also result in use of alternative viral promoters (49) and this interaction is incompletely understood. The reduction in both IE1 and UL138 here may suggest UL4 contributes to establishing the cellular state permissive for appropriate transcriptional transition into reactivation. This is consistent with the current working hypotheses that successful viral reactivation depends on both viral promoter regulation and the coordinated remodeling of host signaling, differentiation, and chromatin regulation (7, 47, 63, 64).

Our data further suggests that UL4 contributes to this process by directly modulating hematopoietic cell biology. Expression of UL4 alone enhanced myeloid colony formation from CD34+ HPCs (**Figure 6A**) and increased both myeloid and erythroid lineage differentiation (**Figure 6B**) and HPC proliferation (**Figure 6C**) indicating when expressed endogenously that UL4 directly influences hematopoietic differentiation programs. Unexpectedly we found that recombinant UL4 failed to reproduce the differentiation phenotype observed with endogenous expression (**Figure 6D-F**). Similarly, conditioned cultures containing UL4-expressing cells exhibited altered cytokine production (Figure 7A and **B**) but did not induce global differentiation of progenitors (**Figure 7C**). Together, these observations suggest that UL4 does not function solely as a conventional soluble ligand acting through paracrine signaling. Instead, UL4 require endogenous expression, specific intracellular trafficking and/or secretion, or interaction with additional viral or cellular transcriptional programs or proteins to exert its specific biological effects. Alternatively, UL4 may initiate cell-autonomous signaling pathways that secondarily remodel the hematopoietic microenvironment. Distinguishing among these possibilities will require identification of cellular binding partners and signaling pathways regulated by UL4 during latent infection.

These findings also extend the concepts that viral gene families can have specific and coordinated regulation of viral latency and reactivation (65, 66) and that immune signaling regulation extends beyond classical immune responses and antiviral activity to latency regulation (7, 50, 63, 67–74). This study also extends the emerging concept that the RL11 gene family may contribute significantly to the viral regulation. Together with previous studies of UL7 (25, 33) and UL8 (27, 75), this work suggests that multiple RL11 family members coordinate the cellular environment required for successful latency and reactivation. Whether these proteins function independently, sequentially, or as components of a integrated regulatory network remains an important question for further study.

This overarching question has several remaining key outstanding gaps. While we demonstrate a clear requirement for UL4 during reactivation, the direct molecular mechanism remains undefined. Similarly, while UL4 expression alters hematopoietic differentiation and cytokine production, both previously linked to regulation of latency and reactivation, whether these activities are directly mechanistically responsible for promoting viral reactivation remain to be fully established. Identification of the UL4-interacting proteins and downstream signaling pathways specific to hematopoietic progenitors will therefore be necessary to determine precisely how endogenous UL4 coordinates that transition from latency to reactivation.

Together, our data suggest a key role for endogenously expressed UL4 as a viral regulator of reactivation in multiple cell types ultimately influencing cellular regulation during the latency to reactivation transition.

## Acknowledgements

This paper is dedicated to Jay A. Nelson. We thank Jay A. Nelson for support of the early studies and critical discussion. We thank Patrizia Caposio for discussion, critical review of the manuscript, and support for early studies including humanized mice. This work was supported by Public Health Services grants: P01 AI127335 to JAN and P01 AI127335 core B to PC for humanized mice from the National Institute of Allergy and Infectious Disease, and P20 GM113126 subproject 6614 to LBC; and startup funding from the University of Nebraska-Lincoln to LBC. CRediT taxonomy: CH: methodology, investigation, data curation, writing – review & editing; KP: methodology, investigation, data curation, writing – review & editing; LBC: conceptualization, methodology, investigation, data curation, validation, formal analysis, writing – original draft, writing – review & edition, visualization, supervision, project administration, funding acquisition.

## Materials and Methods

### Cells

Primary CD34^+^ hematopoietic progenitor cells (HPCs) were isolated as previously described (76) or purchased from StemCell Technologies as deidentified samples obtained from human bone marrow. Human embryonic stem cells (NIH approved WA01 and WA09 hESCs) were differentiated into CD34+ HPCs as previously described (77). All HPCs were cultured in hematopoietic infection media (IMDM (Hylone) supplemented with 10% BIT 9500 serum substitute (StemCell Technologies), 50 μM 2-mercaptoethanol, 2nM L-glutamine, 20 ng/mL human low-density lipoprotein (Sigma-Aldrich) as previously described (76, 77). THP-1 (bone marrow monoblast), NHDF (normal human dermal fibroblast) cells, and 293T cells were obtained from the American Type Culture Collection (ATCC). THP-1 cells were grown in RPMI-1640 (HyClone or Corning) medium supplemented with 10% fetal bovine serum (FBS) (Gibco), 4.5 g/l glucose, L-glutamine, and antibiotics [penicillin (10 units/ml)–streptomycin (10 ug/ml)]. NHDF and 293T cells were cultured in Dulbecco’s modified Eagle’s medium (DMEM) (Hyclone or Corning) supplemented with 10% FBS, penicillin, streptomycin, and L-glutamine (as above). The M2-10B4 murine stromal cell line expressing human interleukin-3 (IL-3) and granulocyte-colony stimulating factor (G-CSF) and the S1/S1 murine stromal cell line expressing human IL-3 and stem cell factor (SCF) were obtained from Stem Cell Technologies and maintained in DMEM with 10% FBS, penicillin, streptomycin, and L-glutamine as above with selection maintained with G418 and Hygromycin (77).

### Viruses

The HCMV TB40/E bacterial artificial chromosome (BAC) was previously engineered to express green fluorescent protein (GFP) as a visual marker of infection(78). Recombinant HCMV TB40/E-UL4stop was constructed by two-step GalK-Kan recombination (79) using pC255 as a template to generate the GalK-Kan insert and the primers HCMV-UL4-GK-F forward (5’-GGCACTGTTTGAGCATGACTGTTTCCAAACCGTAACGTGGTAAATAAATCAGCTCGGTACCCGGGGATCTT-3’) and HCMV-UL4-GK-R reverse (5’-GACGCTTAACATTTCAAAAGCAACGTAAACAATAGCTACAGCCACGCGGCTCTGTGGAACTTCACGTGCGTTCCTCAGCAAAAGTTCGATTTATTCAAC-3’). Following selection of recombinant BACs on the basis of kanamycin resistance, the Kan^R^ marker was removed using minimal DOG selection. Recombination was assessed at each step using flanking primers: HCMV-UL4-flanking-F forward (5’-gtagcgtatccggtttggaa-3’) and HCMV-UL4-flanking-R reverse (5’-tgtcgtaacagcaggattcg-3’) at each step for the removal of wild-type sequence and insertion of GK-Kan insert and removal of the GK-Kan insert and insertion of the UL4stop region, respectively. BAC integrity was examined by enzyme digest analysis and sequencing of the entire viral genome by next generation sequencing. Virus was reconstituted by transfecting the BAC genomes (1 µg) into NHDF fibroblasts and incubating until >80% cytopathic effects (CPE) were observed. Plaque purification was performed and HCMV TB40/E-GFP-UL4stop stocks and titers were generated as previously described (80). UL4stop mutation was assessed by periodic viral genome sequencing to confirm retention of the mutation on later viral stock passages and was confirmed to be maintained through at least passage 7 (virus used between passages 4 and 6 for all experiments). Viruses were titered in parallel (UL4stop and WT) by plaque assay for all experiments.

### Adenovirus Production

Recombinant Adenoviruses were produced by pAdTet7 UL4HA or GFP or empty vector co-transfection of 293 cells with Ad DNA (Ad5-ψ5), as previously described (80) or by using the pAd-Easy system as previously described (81). Recombinant adenoviruses were expanded on 293 cells and the bulk stocks were titered by TCID50 on 293 cells.

### Reagents

Recombinant UL4 protein from TB40E (TB40E: EF999921) was generated at RayBiotech as previously described for UL7 (25)or for a second independent batch by Genscript. Purified, eluted proteins were analyzed by SDS-PAGE and Coomassie blue staining to confirm size. The concentration of purified protein was measured in triplicates using BCA protein assay kit (Thermo/Pierce) with BSA standard. Recombinant IgG Fc was generated and purchased from Genscript.

### Immunoblotting

Extracts were run on 8% SDS-PAGE, transferred to PDVF membrane and visualized with antibodies specific for tag () and anti-rabbit HRP (Cell Signaling Technologies). Total protein staining was performed with the Licor Total Protein stain kit following the manufacturer’s instructions following transfer and before antibody staining.

### Latency and Reactivation

CD34^+^ HPCs were infected at an MOI of 3 for 42 h. Following infection, pure populations of viable, infected (GFP-positive) CD34^+^ HPCs were isolated by fluorescence-activated cell sorting (FACS) using an Aria II (Becton Dickson) using PI for viability and an APC-conjugated CD34 antibody (Biolegend). Long-term cultures were maintained in Myelocult supplemented with hydrocortisone in transwells above an irradiated M2-10B4 and S1/S1 stromal cell monolayer for 12 days. The frequency of infectious center production during the culture period was measured using a limiting dilution assay (82). HCMV latency and reactivation protocols are detailed in depth in (77).

### CD34+ HPC Myeloid Differentiation and HPC Proliferation

Purified CD34^+^ HPCs, infected as above, were seeded at 500 cells/mL (primary), 2000cells/mL (primary bone marrow derived) or 104 cells/mL (hESC-derived) in methylcellulose medium, Methocult H4434 (all primary cells) or Methocult SF (hESC-derived) (StemCell Technologies) at 3 replicate dishes per group. Hematopoietic colonies were scored microscopically 7, 14, and 21 days later using the assistance of a StemVision System (StemCell Technologies).

### Engraftment and infection of humanized mice

All animal studies were carried out in strict accordance with the recommendations of the American Association for Accreditation of Laboratory Animal Care (AAALAC). The protocol was approved by the Institutional Animal Care and Use Committee (protocol 3498) at Oregon Health and Science University. All studies were performed by LBC with assistance from CH and PC. NOD-*scid* IL2Rγ_c_^null^ mice were maintained at a pathogen free facility in accordance with procedures approved by the Institutional Animal Care and Use Committee. Humanized mice were generated as previously described (76). Briefly, huNSG mice were generated by sublethal irradiation following by IV injection of CD34+ HPCs between days 1-3 post-birth. Humanization was determined by flow cytometry analysis every 4 weeks beginning at week 12 post-humanization as previously described (76). Experimental groups were normalized for total humanization (percentage of huCD45+ cells out of total leukocytes (human + mouse CD45+ cells in the periphery) (Supplemental Figure 3A) and for immune cell balance (by percentage of huCD3+ cells out of total huCD45+ leukocytes in the periphery) (Supplemental Figure 3B). Engrafted huNSG mice were then treated with 1mL of 4% Thioglycollate (Brewer’s Media, BD) by intraperitoneal (IP) injection to recruit monocyte/macrophages. At 24hrs post-treatment, mice were infected with HCMV TB40E wild-type or HCMV-UL4stop-infected fibroblasts (approximately 10^5^ PFU of cell-associated virus per mouse) via IP injection. A control group of engrafted mice were mock infected using uninfected fibroblasts. At 8 weeks post infection, the infected mice were split into two groups and half of the mice were treated with 100 1L of G-CSF (Neupogen, at 300 ug/mL; Amgen) using a subcutaneous micro-osmotic pump (1007D; Alzet) and 125 μg AMD3100 (1,1’-[1,4-Phenylenebis(methylene)]bis-1,4,8,11-tetraazacyclotetradecane octahydrochloride or Plerixafor) administered IP to mobilize HPCs. The remaining half of the mice served as a direct comparison for the effects of virus reactivation and dissemination following HPC mobilization. At one week post mobilization, the mice were sacrificed, lymphoid organs harvested for analysis and samples for DNA extraction frozen in RNALater and stored at −80°C for subsequent analysis.

### Quantitative (q) PCR for Viral Genomes and qRT-PCR for Viral Gene Expression

Total DNA from mouse tissues was extracted using the DNAzol kit (Life Technologies), and processed as previously described (83, 84). Total DNA and RNA were extracted from cells using TRIZOL manufacturer’s directions. Primers and a probe recognizing HCMV UL141 were used to quantify HCMV genomes (probe = CGAGGGAGAGCAAGTT; forward primer = 5’ GATGTGGGCCGAGAATTATGA and reverse primer = 5’ ATGGGCCAGGAGTGTGTCA) as previously described. Viral genomes in humanized mice were normalized to 11g input DNA. Viral genomes synthesized during infection in CD34^+^ HPCs were normalized to total cell number determined using human β-globin as a reference (probe = GGACAGATCCCCAAAGGACT; forward primer = 5’ TTAGGGTTGCCCATAACAGC and reverse primer = 5’ TTGGACCCAGAGGTTCTTTG) as previously described using TaqMan FastAdvanced. IE1 and UL138 were detected using SybrGreen qRT-PCR (Abcam) and previously published primers (85).

### Statistical Analysis

Statistical analysis was performed using GraphPad Prism (v9) for comparison between groups using an unpaired Student’s t-test or one-way Anova or two-way Anova with p values as indicated in each figure.

---

**Supplemental Figure 1.**
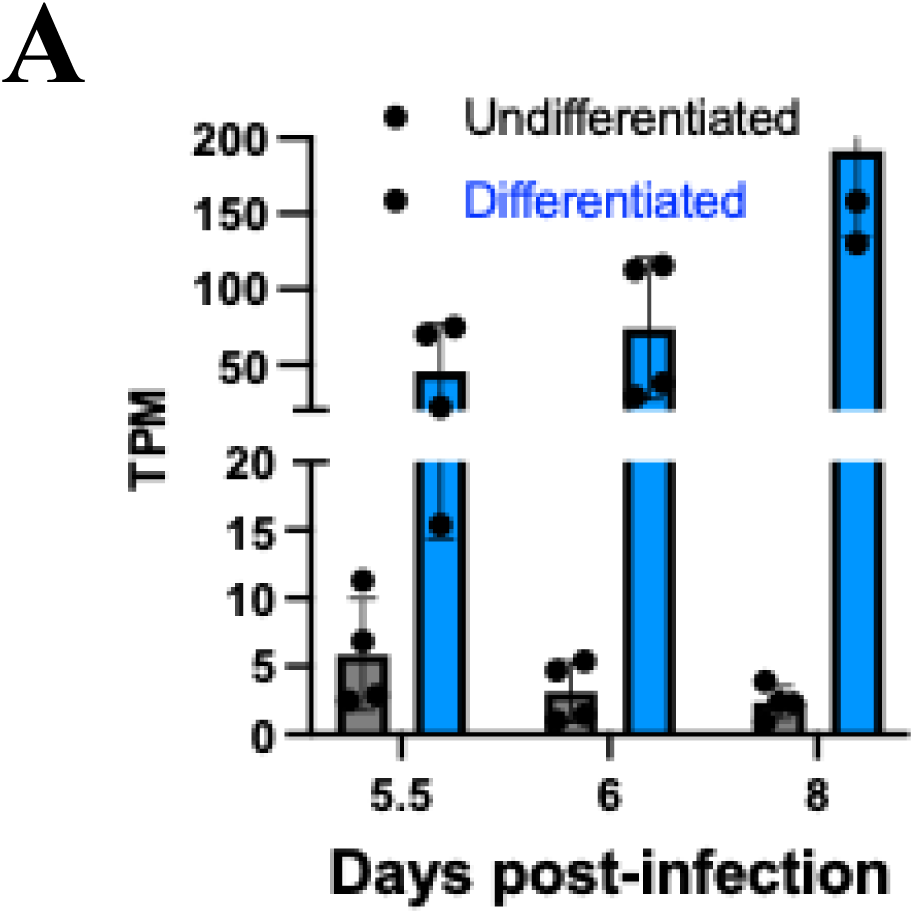
Expression of UL4 in a THP-1 model of latency and reactivation. Raw data from the Goodrum lab, published in: Collins-McMillen et. al., 2025. PMID: 40434638 (47).

**Supplemental Figure 2.**
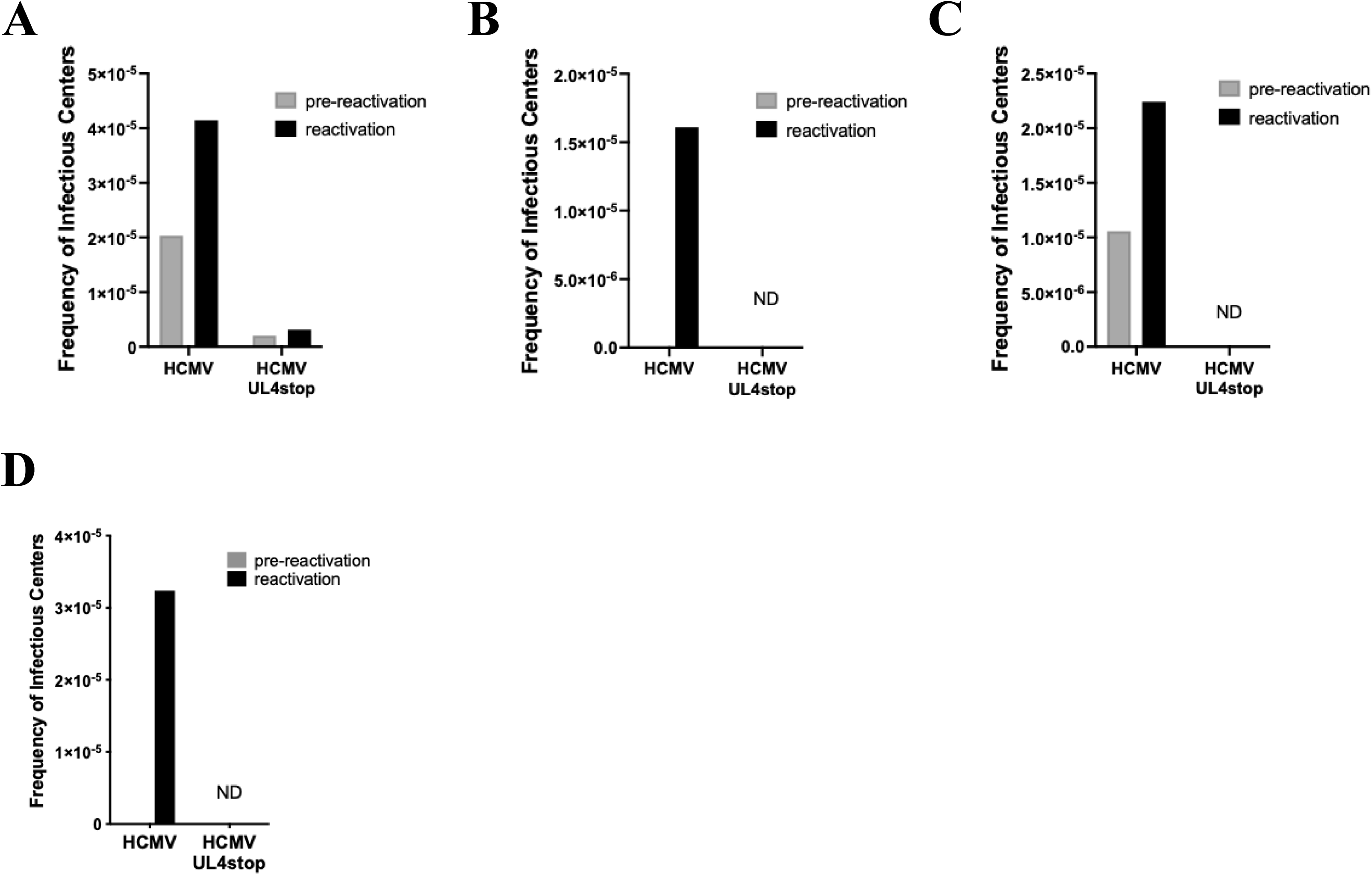
Individual HPC donors from Figure 3: UL4 is required for latency and reactivation in CD34+ HPCs. CD34+ HPCs infected with HCMV or HCMV lacking UL4 (HCMV-UL4stop), were sorted, and cultured for latency as in Figure 2. At 14dpi (12d latent), equivalent populations of cells were either directly plated on fibroblasts in the presence of cytokine support (reactivation) or lysed and plated on fibroblasts to measure the amount of virus present during latency (pre-reactivation), and the frequency of infectious centers calculated by ELDA. Data shown is the frequency of infectious center formation for each of 4 independent primary HPC donors (A through D).

**Supplemental Figure 3.**
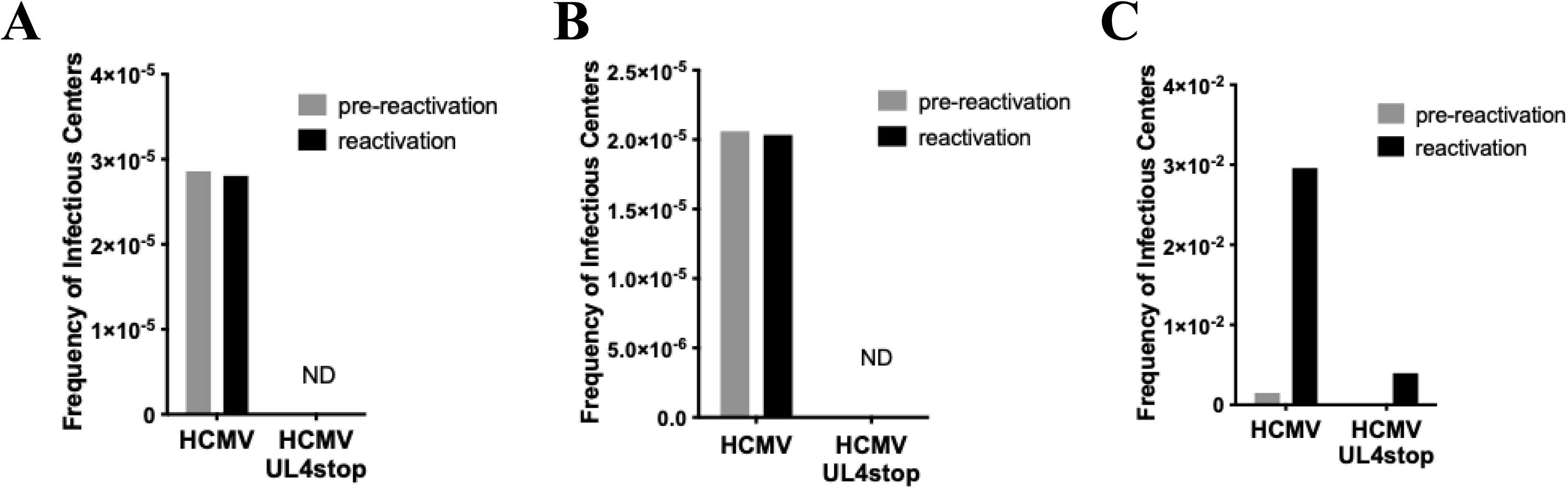
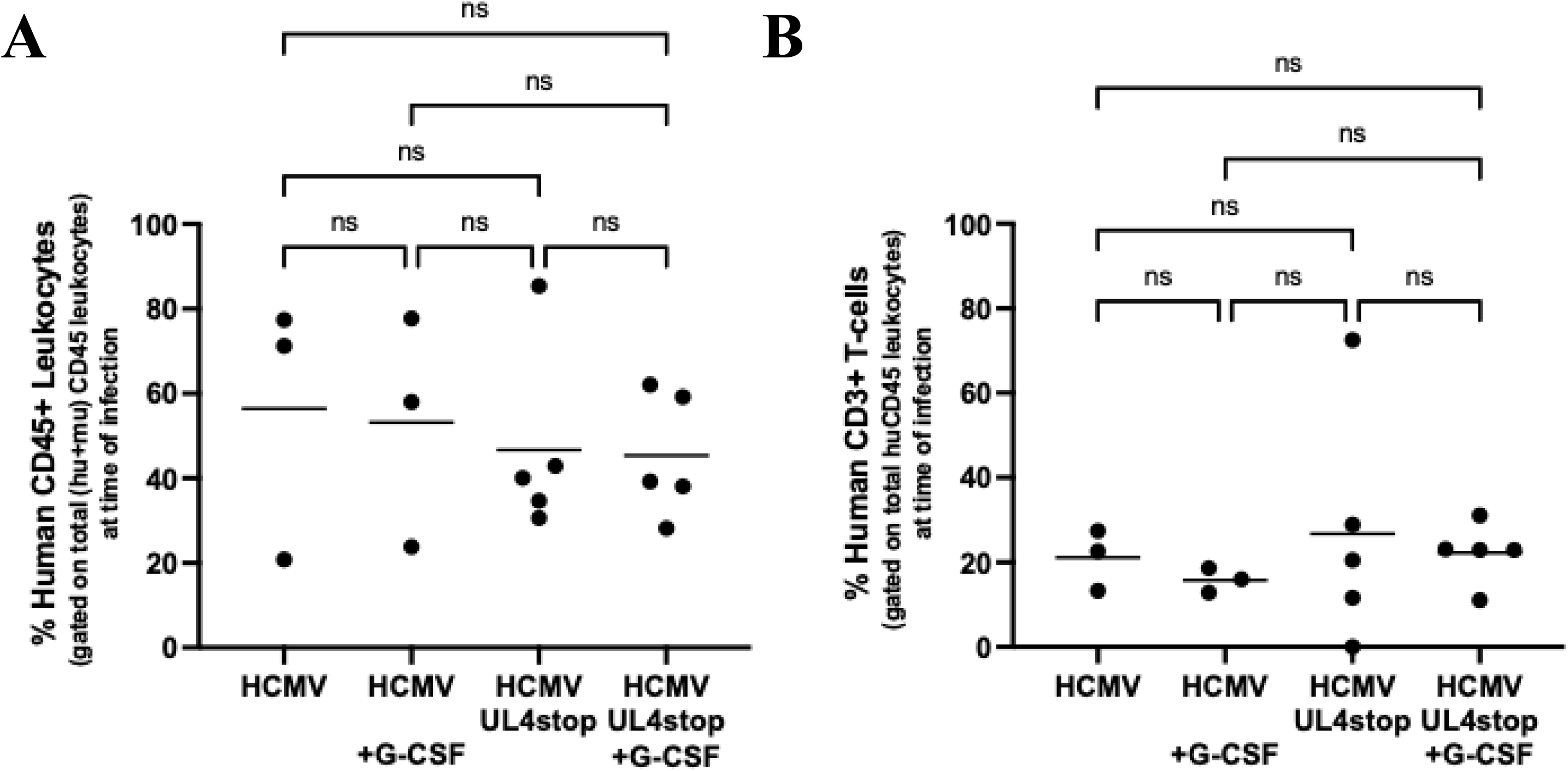
Humanization Parameters of huNSG mice used in Figure 5 prior to infection. huNSG mice were generated by intrahepatic injection of CD34+ HPCs into sublethally-irradiated neonatal mice as described. Beginning at 12 weeks post-humanization, mice were screened by peripheral blood draw and flow cytometry for humanization. Blood samples were lysed for red blood cells, washed, stained with a fixable viability dye, blocked in human and mouse serum, and stained with antibodies specific to human and mouse cell surface markers for phenotyping. Data shown is from the blood draw immediately preceding infection when groups were normalized for engraftment (**A**) and immune cell populations (**B**). **A)** Total human CD45+ cells as a percentage of human and mouse CD45+ leukocytes (to normalize differences in blood draw volumes). **B)** Human CD3+ T-cells as a percentage of total human CD45+ cells. Data shown is for all individual mice graphed per their final viral group assignment. NS = not significant by ANOVA.

**Supplemental Figure 4.**
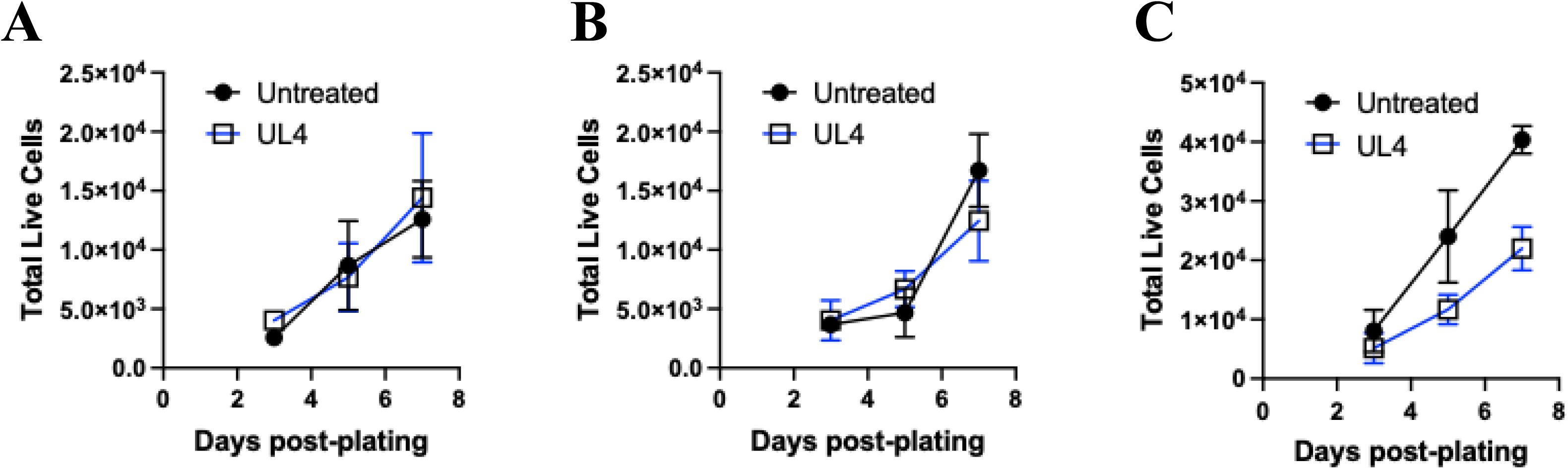
Replication Experiments for Figure 6. Endogenous but not exogenous UL4 controls HPC functional outcomes. HPCs were treated with exogenous UL4 protein and plated in cytokine enhanced liquid media for proliferation Data shown in A, B, C, are 3 independent experiments with different human cell donors. *p<0.05, NS = not significant.

## Notes

### Competing Interest Statement

The authors have declared no competing interest.

